# SEMA3F-deficient colorectal cancer cells Promote lymphangiogenesis: fatty acid metabolism replace glycolysis for energy supply during lymphatic endothelial cells proliferation in tumor hypoxia microenvironment

**DOI:** 10.1101/869644

**Authors:** Xiaoyuan Fu, Miaomiao Tao, Hongbo Ma, Cancan Wang, Yanyan Li, Xiaoqiao Hu, Xiurong Qin, Renming Lv, Gengdou Zhou, Jun Wang, Meiyu Zhou, Guofa Xu, Zexin Wang, Min Chen, Qi Zhou

## Abstract

lymphangiogenesis as a process is colorectal cancer first metastasis via lymphatic vessels to proximal lymph nodes. The fuel metabolism in mitochondrial and support proliferation of lymphatic endothelial cells (LECs) remain elusive during lymphangiogenesis in tumor hypoxic microenvironment. Recent studies report that loss of SEMA3F critically contributes to lymphangiogenesis of the CRCs. Here, we silenced SEMA3F expression of CRCs and co-culture with hLECs, the tubulogenesis capacity and hLECs migration were escalated in the hypoxia, the hLECs mainly relied on fatty acid metabolism not aerobic glycolysis during lymphangiogenesis. SEMA3F-deficient CRCs up-regulated PMAKP expression and phosphorylation of hLECs, and activated its peroxisome proliferator-activated receptor (PPARs) and Peroxisome proliferator–activated receptor gamma coactivator-1 alpha (PGC-1a) facilitated their switched toward fatty acids (FA) catabolism. Furthermore, we observed that activation of the PGCI-PPAR lipid oxidation signaling pathway in hLECs was caused by the secretion of interleukin-6 by tumor cells.Taken together, this study indicates that CRCs with SEMA3F expression depletion significantly promotes lymphangiogenesis in hypoxia and faciliates *the* secretion *of* IL-6 in tumor cell, and activates mitochondria fatty acids oxidation (FAO) reaction in the hLECs by PGCI-PPAR signaling pathways to support its growth.

## Introduction

The growth and metastatic of colorectal adenocarcinoma (CRC) often are the leading cause of mortality in patients, colorectal cancers metastasis lead spread of cancer cells, where cancer cells often disseminate through blood and lymphatic vessels (Hanahan and Weinberg, 2011; Ma *et al*,2018). Lymphatic invasion is one of the most events and lymph node involvement represents one of the most important prognostic factors of poor clinical outcome in patients with CRC (Royston and Jackson, 2009; Sakurai *et al*,2012; Veronese *et al*,2016; Vellinga *et al*,2017). However, the role of lymphangiogenesis and the molecular mechanisms underlying lymphangiogenesis in cancer remains largely elusive, such as interactions among lymphatic endothelial cells, tumor cells, and other components within the tumor microenvironment (TME). Specifically, the common features in solid tumors TME is hypoxia and acidosis, the lymphatic endothelial cells (LECs) energy metabolism and it’s proliferation are far from full understand. Although the molecular mechanisms angiogenesis is required for solid tumors is well known. Class III semaphorins is involved in tumor angiogenesis, lymphatic vessels formation and axonal guidance regulation (Tessier-Lavigne and Goodman,1996; Tran *et al*,2009). semaphorin-3F (SEMA3F) is a member of the semaphorin-3 family(Chen *et al*,1998;Bielenberg *et al*,2004; Fu and Ip,2017), This secreted semaphorins, acting via binding neuropilins (NRPs), Chemorepulsive for vascular endothelial cells and lymphatic endothelial cells expressing Neuropilin-2 (NRP2) (Bielenberg *et al*,2004; Doçi *et al*,2015). SEMA3F has higher binding affinity for NRP2, the recombinant SEMA3F promotes LECs collapse and potently inhibits lymphangiogenesis in vivo, and the SEMA3F re-expression diminishes lymphangiogenesis and lymph node metastasis in vitro (He and Tessier-Lavigne, 1997; Bielenberg *et al*,2004; Klagsbrun and Shimizu, 2010; Sakurai *et al*,2012). In our previous study, loss of SEMA3F, the inhibitory ligand of NRP2, critically contributes to CRCs metastasis (Wu *et al*, 2011). Furthermore, our previous study has demonstrated that NRP2 significantly increased CRCs lymphangiogenesis and induced the activation of NRP2 in hLECs to promote tumor lymphangiogenesis via integrinα9β1/FAK/Erk pathway independent VEGF-C/VEGFR3 signaling (Ou *et al*, 2015). It suggest that SEMA3F maybe an anti-lymphangiogenic metastasis suppressor during cancer progression. Intriguingly, The class 3 semaphorins was initially identified as axonal repellents and acted via NRPs, suppressed axons from growing sensory, sympathetic, and motor neurons (Lumb *et al*, 2018; Inokuchi *et al*, 2017; Takeuchi *et al*, 2010). In early studies, SEMA3F via Neuropilin-2, to repulsive sensory and sympathetic axon repulsion, principally responsible for mediating responses SEMA3F was Plexin-A3 (Coate *et al*, 2015; Wang *et al*, 2017). Recent evidence pointed to that the sympathetic nervous system (SNS) could control brown adipose tissue function and energy homeostasis (Nakamura *et al*, 2017; Riera *et al*, 2017; Owen *et al*, 2014; Morrison *et al*, 2014). It suggest sympathetic nerve implicate lipid metabolism, it also suggest SEMA3F maybe has potential role in energy availability. The other studies demonstrate the epigenetics may control differentiation and fatty acid metabolism of LECs ( Wong *et al*, 2017; van der Klaauw *et al*, 2019). So, We hypothesize that SEMA3F binding NRP2 to reduce lymphangiogenesis of CRCs maybe has other mechanisms, SEMA3F may bind NRP2 to disturb lipid metabolism of LECs and repress lymphangiogenesis of CRCs.

In this study, we examined the role of SEMA3F expression deficient CRCs promotion hLECs lipid metabolism and lymphangiogenesis under hypoxemia.We demonstrated that SEMA3F plays a crucial role in hLECs energy metabolism. CRCs with SEMA3F expression depletion significantly promotes hLECs migration and tubulogenesis capacity in hypoxia, and enhance the lipid metabolism via suppressing aerobic glycolysis of hLECs, and the lipid metabolism to induce the PGCI-PPAR signaling pathway in hLECs. Further evidence demonstrated that CRCs cell with SEMA3F expression depletion up-regulated its IL-6 secretion, to activate the PGCI-PPAR signaling pathway and induces the lipid metabolismin of hLECs in hypoxia. Our findings for the first time revealed a novel role and the underlying mechanism, the CRCs cell with SEMA3F expression deficiency was involved the energy metabolism with hLECs in the hypoxia environment, and induce tumor lymphangiogenesis of CRCs to promote lymphatic metastasis.

## Result

### 1. SEMA 3F-deficient CRCs further enhance lymphangiogenesis under hypoxic conditions

In our previous study, loss of SEMA3F expression, the inhibitory ligand of NRP2, critically contributes to CRCs metastasis and lymphangiogenesis in normoxia(Wu *et al*, 2011; Ou *et al*, 2015). To examine whether SEMA3F loss expression plays any roles in induce lymphangiogenesis of colorectal cancer cell in hypoxia, We used supernatants of the colorectal adenocarcinoma cell line HCT116 with SEMA3F expression and SEMA3F knockout (SEMA3F KD), to culture hLECs three-dimensionally on matrigel substrate under normoxia and hypoxia to induce tubule formation of hLECs. The tubulogenesis of hLECs was improved in hypoxia than that in normoxia (p < 0.001; Figure 1A and B). Strikingly, comparing with control cells, SEMA3F KD groups displayed significantly enhancement of tubulogenesis ability of hLECs under normoxic conditions. Consistent with normoxic conditions culture, SEMA3F KD groups had stronger ability to induce hLECs tubule formation than SEMA3F expression groups in hypoxia. We used a co-cultured system consisting both hLECs and CRCs with transwell chambers respectively in normoxic and hypoxic conditions culture to induce migration of hLECs. It had been shown that (Figure 1C) SEMA3F KD groups promoted migration of hLECs in normoxia, futhermore, SEMA3F KD groups further enhanced capability of hLECs migration in hypoxia. These effects indicated loss of SEMA3F, CRCs further promoted lymphangiogenesis in hypoxia.

**Figure 1.**
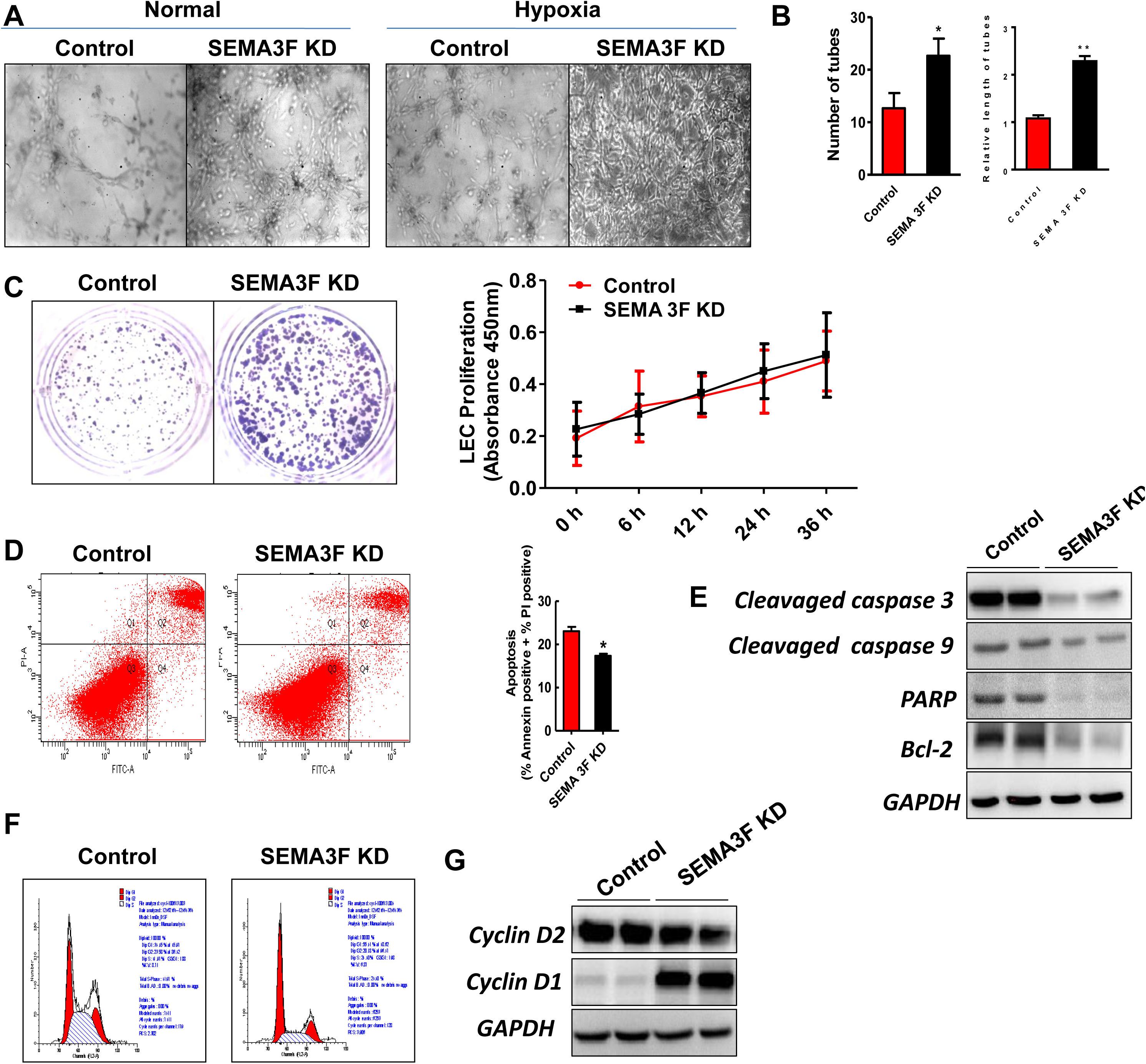
SEMA3F deficient further contribute to CRCs cell-induced migration and tubulogenesis of hLECs at the hypoxia condition. A, SEMA3F knockdown in CRCs enhance tubulogenesis of hLECs both normoxia or hypoxic, which is more pronounced at hypoxia. B, Hypoxia culture in vitro, SEMA3F knockdown of CRCs increased the number and length of tubules of hLECs(*p<0.05, **p<0.01). C, SEMA3F knockdown in CRCs promote hLECs migration using transwell assay and proliferation at hypoxia than those in in normoxia. D, The CRCs knocked down by SEMA3F during hypoxia significantly reduced the apoptosis of hLECs using flow cytometry of Annexin V/PI (Sigma) staining compared with the normoxic group(*p<0.05). E, Western blot analysis confirmed that SEMA3F knockdown in CRCs resulted in decreased expression of hLECs pro-apoptotic and anti-apoptosis-related gene proteins, and the hypoxic culture group was more markedly reduced than the normoxic culture group. F, Compared with the normoxic group, flow cytometry detection of SEMA3F knockdown in CRCs significantly increased the proportion of G1 phase of hypoxic cultured hLECs and promoted the proliferation of hLECs. G, Western blotting determined that SEMA3F knockdown in CRCs could increase the expression of cyclinD1 in hLECs cultured in hypoxia.

To gain further cause insight about promotion of lymphangiogenesis, we further confirmed with flow cytometry assessment that the apoptosis rate of hLECs in SEMA3F KD groups was lower than it of control groups in hypoxia (p < 0.005; Figure 1D). We then measured apoptosis and proliferation relative gene expression levels of hLECs, it showed the expression level of pro-apoptotic protein Caspase3 and 9,and the main shearing object poly ADP-ribose polymerase (PARP) of caspase 3 was decreased (Figure 1E) in SEMA3F KD groups, but the anti-apoptotic protein BCL-2 still was decreased. However, the expression level reduction in the SEMA3F KD groups was more pronounced than in the control groups, which may be the reason for the lower apoptotic rate in the SEMA3F KD groups. In addition, the expression of cyclinD1 was up-regulated in SEMA3F KD groups under hypoxic condition, which significantly increased the proportion of G1 phase hLECs and promoted the proliferation of hLECs(Figure 1F and G), not cyclinD2. Thses effects revealed SEMA3F deficiency in CRCs could further enhance lymphangiogenesis in the hypoxia compared to the normoxic conditions via promotion hLECs proliferation, not influence its apoptosis.

### 2. SEMA 3F-deficient CRCs promote lipid peroxidation of mitochondria in LECs at the hypoxic conditions

To explored causal of energy supply of hLECs proliferation when we silenced SEMA 3F expression in CRCs, we employed the CRCs which was control groups and respectively co-culture with hLECs in hypoxia. Interestingly, we found that the triglycerides level in hLECs which co-culture with SEMA3F KD-CRCs groups was three times higher than control groups, but the free cholesterol, total cholesterol and phospholipids level were no difference between the two hLECs groups (P<0.05;Figure 2A), it showed SEMA3F KD-CRCs groups increased hLECs cellular content of triglycerides in hypoxia. We presumed that the energy supply that SEMA3F KD-CRCs could induce hLECs proliferation under hypoxia was derived from fat metabolism. Thus, we detected the protein expression level of fat metabolism enzyme in two hLECs groups with western blot. As expected, the triglyceride synthesis genes such as glycerol-3-phosphate acyltransferase (GPAT) and phospholipase C(PLC) expression level of hLECs co-culture with SEMA3F KD-CRCs groups were increased obviously than control groups at hypoxia(Figure 2B left). These data demonstrated that SEMA3F KD-CRCs enhanced hLECs triglyceride synthesis and acted on first committed step of their synthesis, it is the acylation of GPAT enzymes via the glycerol phosphate pathwayphosphatidylinositol (PI)-specific phospholipase C (PLC), which reside in the endoplasmic reticulum (ER) and mitochondria (Cao et al,2006; Ohba et al,2013; Nagarajan *et al*,2017; Tang *et al*,2017). Consistent with results of immunofluorescence examination showed that the activity of mitochondria in hLECs which co-culture with SEMA3F KD-CRCs groups was higher than control group (p<0.001;Figure 2C and F).

**Figure 2.**
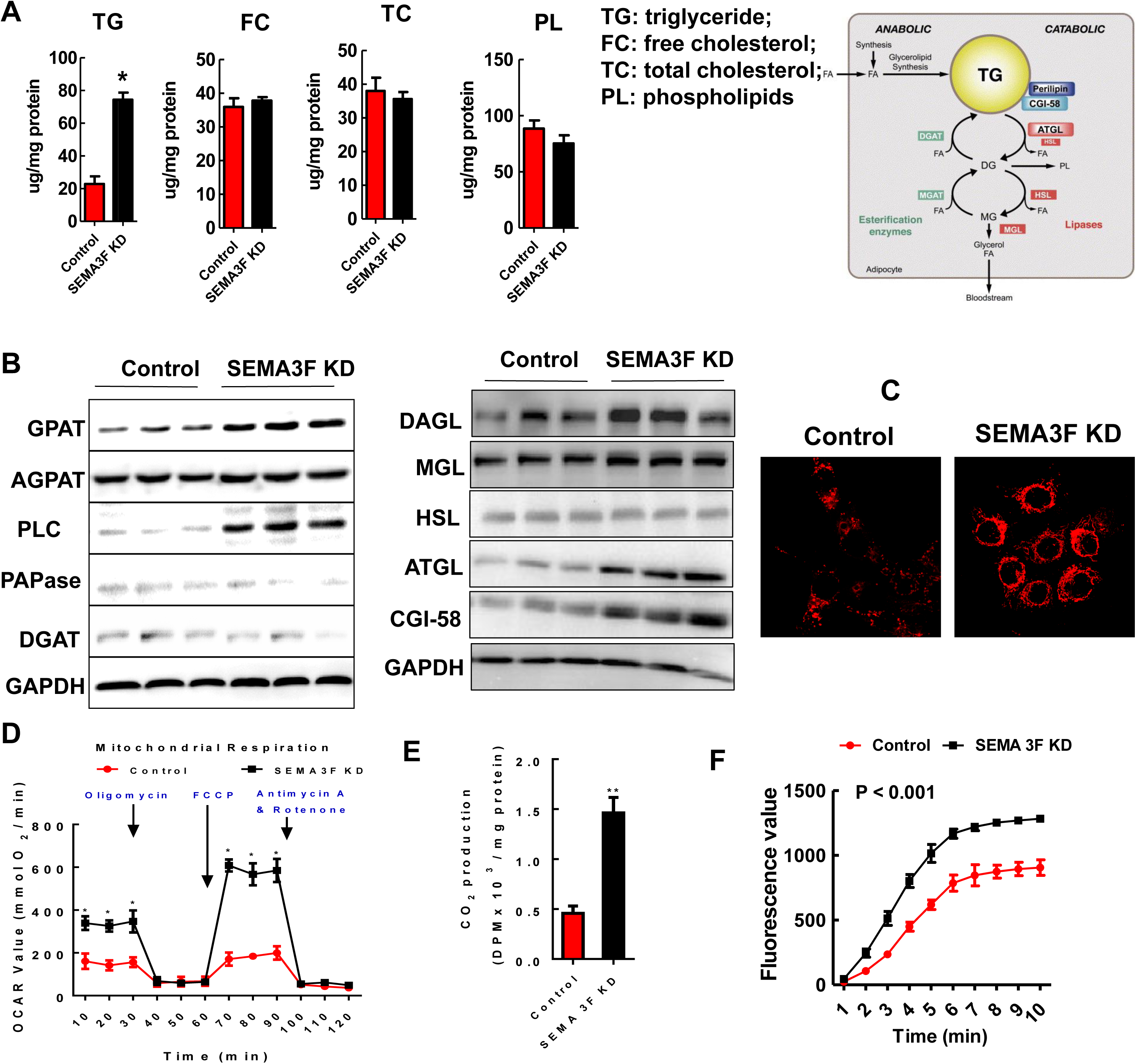
SEMA 3F-deficient CRCs under hypoxic conditions promote mitochondrial lipid oxidation in hLECs. A, The triglyceride levels of hLECs co-cultured with SEMA 3F-deficient CRCs under hypoxic conditions were elevated, but free cholesterol, total cholesterol and phospholipid levels did not differ between the normoxic and hypoxic culture groups(*p<0.05). B, Western blot analysis of the protein levels of GPAT, AGPAT, PLC, PAPase, DGAT, DAGL, MGL, HSL, ATGL, CGI-58 in hLECs co-cultured with SEMA 3F-deficient CRCs and control group at hypoxia condition. C, Immunofluorescence showed the activity of mitochondria in hLECs which co-culture with CRCs in SEMA3F knockdown and control group at hypoxia condition. D, The mitochondrial respiratory chain complex Oligomycin, Fccp and Antimycin A & Rotenone level in hLECs which co-culture with CRCs in SEMA3F KD groups and control group at hypoxia condition(*p<0.05). E, The concentration of CO2 productions of hLECs which co-culture with CRCs in SEMA3F KD groups and control group at hypoxia condition(**p<0.01). F, the mitochondrial fluorescence value of hLECs which co-culture with CRCs in SEMA3F KD groups and control group at hypoxia condition(p<0.01).

We then tested situation of hLECs lipolysis, which co-culture with SEMA3F KD groups, to explore it energy supply in hypoxia. Shown in Figure 2B right, the catabolism relative genes adipose triglyceride lipase(ATGL),diacylglycerol lipase(DAGL) and comparative gene identification-58(CGI-58) expression level of hLECs was significantly increased in SEMA3F KD groups than those in control groups. These data indicated to knock down SEMA3F of CRCs could promote mitochondria lipid oxidation in hLECs at hypoxia. In order to further clarify the role of SEMA3F KD-CRCs promote mitochondria lipid oxidation in hLECs at hypoxia, we inhibited metabolism of LECs using the inhibitor of three respiratory chain complex oligomycin and antimycin A & rotenone. Our results showed that Oligomycin and Antimycin A & Rotenone level in hLECs which co-culture with SEMA3F KD-CRCs groups were respectively two times and three times comparing with control groups (P<0.05;Figure 2D), however, it was not difference of Fccp between two groups at hypoxia. It indicated the respiratory functions in hLECs which co-culture with SEMA3F KD-CRCs groups was much stronger than control groups. Moreover, the concentration of CO_2_ productions in hLECs which co-culture with SEMA3F KD-CRCs groups was three times higher than control groups (P<0.01; Figure 2E). It indicated that to knock down SEMA3F of CRCs could promote LECs proliferation in hypoxia, it was by promoting their mitochondria lipid oxidation to gain energy.

### 3. SEMA 3F-deficient CRCs suppress glycolysis metabolism of LECs in hypoxia

It is well known that the major energy supply mode of cell is glycolysis when it is in hypoxia.To knock down SEMA3F of CRCs whether can promote glycometabolism of mitochondria in hLECs at hypoxia? The first rate-limiting step in glucose utilization in cells is facilitation glucose transport(Birsoy *et al*,2014;Nagarajan *et al*,2017; Tang *et al*,2017). In present study, we examined the expression of the glucose transporter protein1 (GLUT1) in the hLECs of co-culture with SEMA3F KD-CRCs groups and control groups in hypoxia. The western blotting results showed the GLUT1 protein expression level was reduced in SEMA3F KD groups comparing that in control groups (Figure 3A). We further examined basal glucose uptake and insulin-stimulated glucose uptake of hLECs in two groups. The results showed that the basal glucose uptake and insulin-stimulated glucose uptake in SEMA3F KD groups were markedly lower than those of the controls(P<0.05;Figure 3 B). These results collectivey illustrated the SEMA3F KD in CRCs repressed glucose uptake and transportation in hLECs at hypoxia.

**Figure 3.**
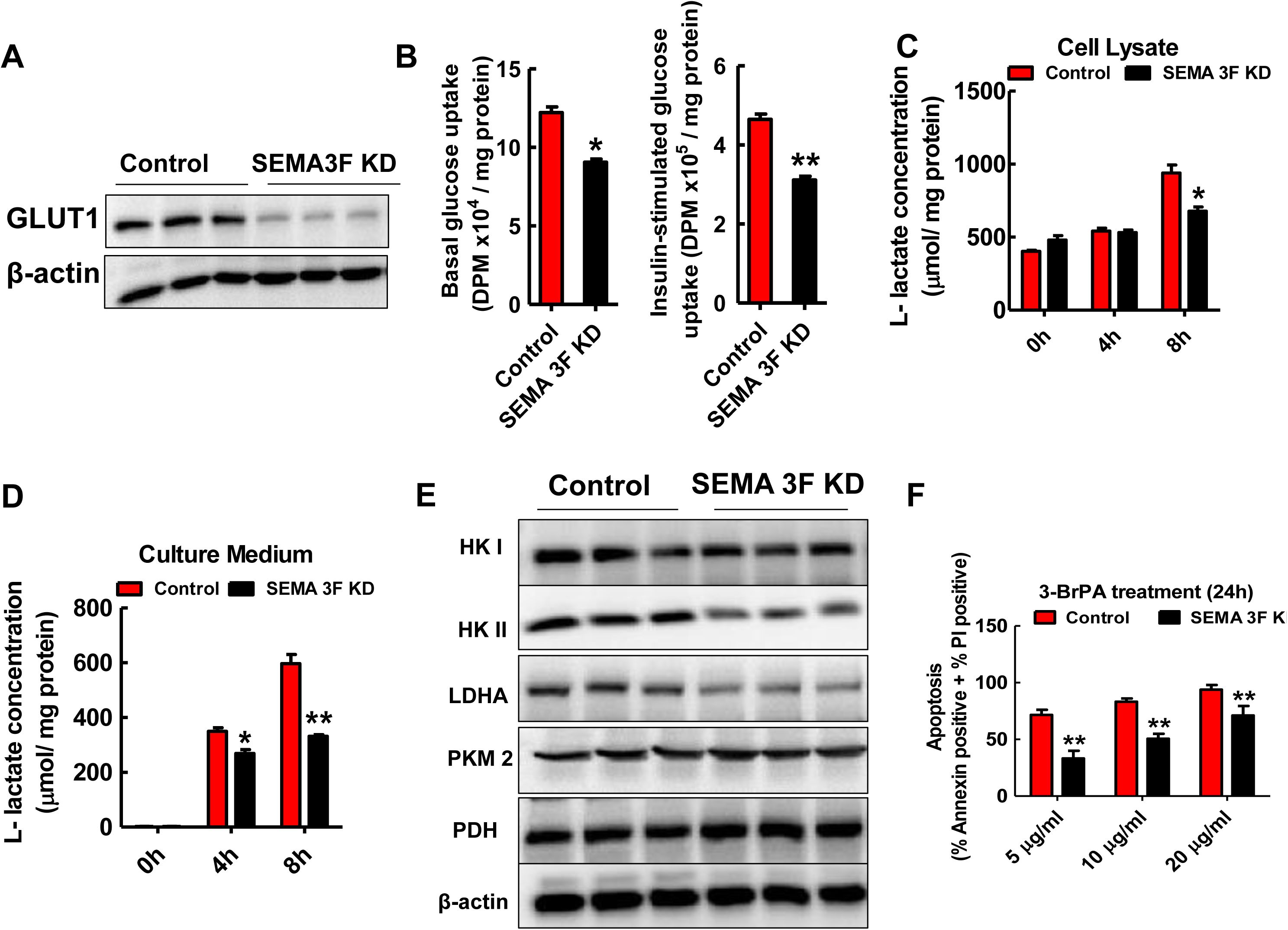
SEMA 3F expression deletion CRCs inhibited glycometabolism of hLECs at the hypoxia condition. A, Western blots analysis the expression levels of GLUT1 of hLECs which co-culture with CRCs in SEMA3F KD groups and control groups at hypoxia condition. B, The basal glucose uptake and insulin-stimulated glucose uptake level of hLECs which co-culture with CRCs in SEMA3F KD groups and control groups at hypoxia condition (*p<0.05, **p<0.01). C, The L- lactate concentration of hLECs lysate which co-culture with CRCs in SEMA3F KD groups and control groups at hypoxia condition(*p<0.05). D, The L- lactate concentration of hLECs culture medium which co-culture with CRCs in SEMA3F KD groups and control groups at hypoxia condition(*p<0.05, **p<0.01). E, Western blots analysis the exprssion levels of HK I, HK II, LDHA, PKM2 and PDH in hLECs which co-culture with CRCs in SEMA3F KD groups and control groups at hypoxia condition. F, The apoptosis rate of hLECs which co-culture with CRCs in SEMA3F KD groups and control groups at hypoxia condition after 3-bromopyruvate treatment(**p<0.01).

To further confirm whether SEMA3F KD in CRCs inhibition glycolysis of hLECs in hypoxia, we respectively tested the L-lactate concentration of the supernatant and hLECs cell lysates in the two groups (Faubert *et al*,2017). As expected, the L-lactate concentration of the supernatant and cell lysate in SEMA3F KD groups were significantly lower than those in control group (p<0.05). Moreover, the differences was much significant with the co-culture time extension (p<0.05;Figure 3C and D). We further tested the genes expression level of glycolysis, it revealed the glycolysis relative genes expression level of hLECs co-culture with SEMA3F KD-CRCs was strikingly lower compareing with control groups in hypoxia (Figure 3 E), such as hexokinase I (HK I), hexokinase II (HK II) and LDHA. In contrast, the pyruvate dehydrogenase levels was higher in SEMA3F KD groups, but Pyruvate kinase isozymes M2(PKM2) expression levels was not difference between two groups. These results suggested that level of aerobic glycolysis of LECs in SEMA3F KD groups was lower than that in control groups in hypoxia, it converted glycolysis to the Krebs cycle was not be blocked(Liu *et al*,2017;DeWaal *et al*,2018; Sukonina *et al*,2019). To further determination the efficacy to knock down SEMA3F of CRCs inhibition glycolysis of hLECs in hypoxia, therefore, we use 3-bromopyruvate to inhibit the glycolysis of the two groups. The result was showed in Figure 3F, the apoptosis rate of hLECs in SEMA3F KD groups were significantly lower than control groups(p<0.01), furthermore, it showed dose-effect relationship. It suggested that, in hypoxia, the energy metabolism in hLECs which co-culture with SEMA3F KD-CRCs not depended on the glycolysis, and also inhibited its glycometabolism. Its major energy supply maybe come from mitochondria lipid oxidation. our data agree with Wong et al report that the fatty acid oxidation (FAO) flux in LECs is even higher than in BECs while glycolytic flux is lower(Wong *et al*, 2017).

### 4. SEMA3F-deficient CRCs promote the activation of lipid oxidation signaling pathway PGCI-PPAR in the hLECs under hypoxia

To confirm the energy metabolism in hLECs which co-culture with SEMA3F KD-CRCs under hypoxia come from mitochondria lipid oxidation, whether the lipid oxidation signaling pathways should be activated. So, we detected the protease expression level which was related to lipid oxidation signaling pathway in two hLECs groups. As shown in Fig.4 A, in hypoxia, adenosine 5’-monophosphate activated protein kinase (AMPK) and phospho-AMPK(P-AMPK), the metabolic sensor or “fuel gauge” (Julien *et al*, 2017), their expression level in hLECs which co-culture with SEMA3F KD-CRCs groups was significantly elevated,but the ratio between p-AMPK /AMPK was decreased. It indicated in the AMP:ATP ratio was increased and ATP-consuming switched to FAO and generated ATP-producing. (Weinberg *et al*, 2015) Furthermore, the expression of lipid oxidation signaling pathway relative proteases also were increased obviously such as PGC1α and PPARα in hLECs co-culture with SEMA3F KD-CRCs groups than those in control groups. However, the PPARγ was not change in two groups (Figure 4A). Simultaneously, PPAR activity of hLECs was also obviously elevated in SEMA3F KD groups (p<0.05; Figure 4B). When we silenced the AMPKα1 or PGC1α expression of hLECs in two groups by siRNA in hypoxia, the PPAR activity of hLECs in control groups has no changes (Figure 4C), conversely, the PPAR activity in hLECs co-culture with SEMA3F KD-CRCs groups was significantly decreasing (p<0.05). In addition, when we silenced the AMPKα1 expression, the hLECs tubulogenesis was significantly decreased at hypoxia in SEMA3F KD-CRCs co-culture groups (p<0.05;Figure 4D). These data showed that hypoxia maybe promote activation of AMPK in hLECs with SEMA3F KD-CRCs co-culture groups, and gain energy via their lipid oxidation signaling pathways in mitochondria. Thus, we measured the change of P-AMPKα and PGC1α protein level in hLECs co-culture with SEMA3F KD-CRCs groups after adding PPAR γ antagonist at the hypoxia. After adding PPAR γ antagonist, the P-AMPKα and PGC1α expression of hLECs both decreased in SEMA3F KD groups, (Figure 4E). These data suggested that in hypoxia, SEMA3F KD in CRCs enhanced hLECs mitochondria oxidation reaction via PGCI-PPAR signaling pathways.

**Figure 4.**
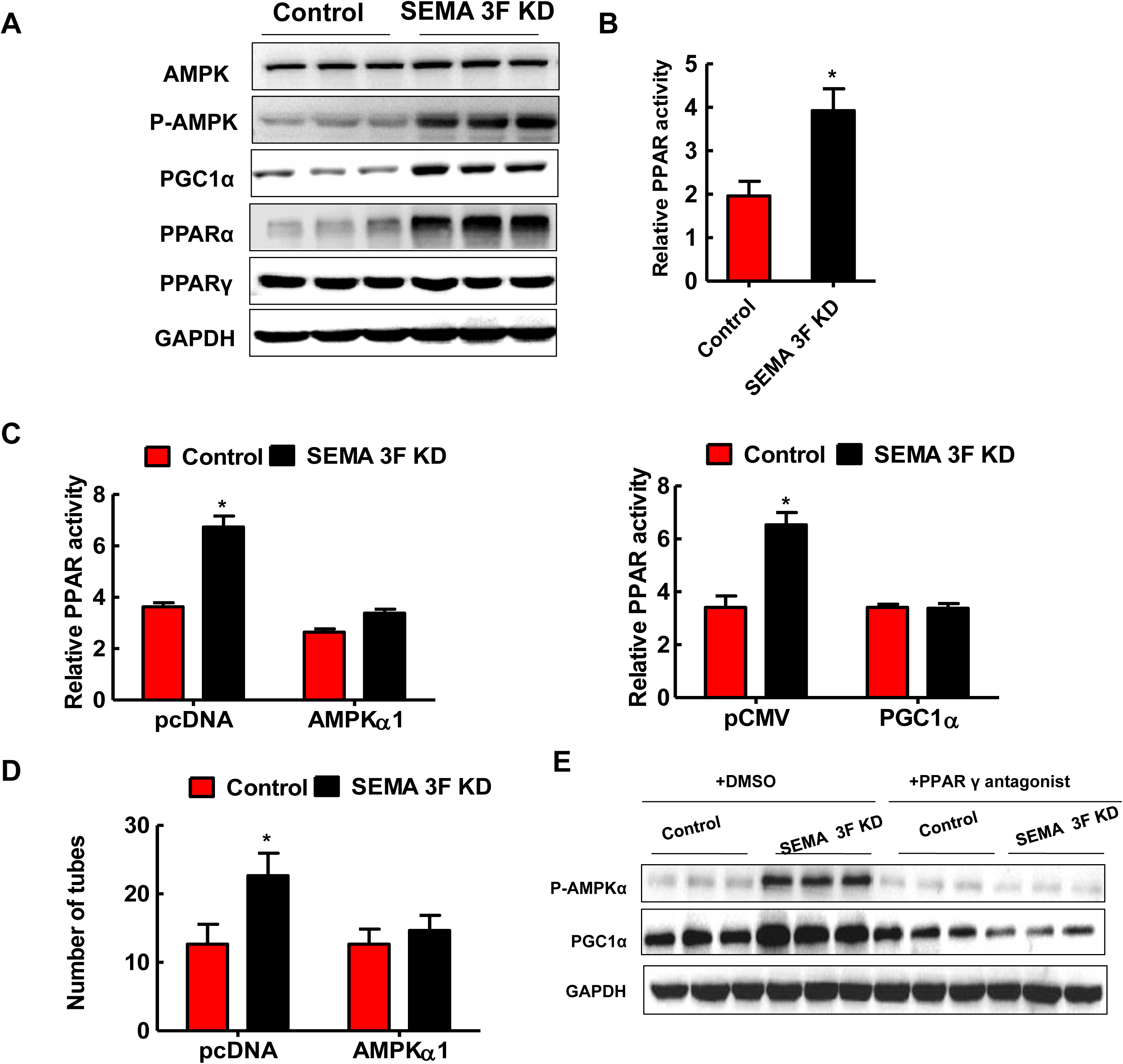
CRCs with SEMA 3F KD promoted the activation of lipid oxidation signaling pathway PGCI-PPAR in hLECs at the hypoxia condition. A, Lysates from hLECs which co-culture with CRCs in SEMA3F KD groups and control groups were immunoblotted for AMPK,P-AMPK,PGC1α,PPAR, PPARγ at hypoxia condition. B, The PPAR activity of hLECs which co-culture with CRCs in SEMA3F KD groups and control groups at hypoxia condition(*p<0.05). C, Comparison of the PPAR activity of LECs without and with AMPKα1 or PGC1α KD, which co-culture with CRCs in SEMA3F KD groups and control groups at hypoxia condition(*p<0.05). D, Comparison of the total tubes number sprout from hLECs without and with AMPKα1KD, which co-culture with CRCs in SEMA3F KD groups and control groups in hypoxia (*p<0.05). E, Lysates from hLECs with and without PPAR γ antagonist which co-culture with CRCs in SEMA3F KD groups and control groups in hypoxia were immunoblotted for P-AMPKα, PGC1α.

### 5. SEMA3F-deficient CRCs enhance IL-6 secretion to drive the lipid oxidation activation of PGCI-PPAR signaling pathway in LECs in hypoxia

Previous research has shown that IL-6 could promote the lipid oxidation by raising PPARα in murine models of fatty liver (Hong *et al*, 2004; Awazawa *et al*, 2011). Whether SEMA3F KD in CRCs initiate the mitochondrion lipid oxidation in hLECs also by enhancing the its secretion of IL-6 in hypoxia? Therefore, we tested the IL-6mRNA level of CRCs in SEMA3F KD groups when it was culture in hypoxia, it was much higher than control groups after culture for 24h (p<0.05;Figure 5A). In order to confirm SEMA3F KD promote IL-6 secretion of CRCs at the hypoxia condition, we further tested the IL-6 level in supernatant of CRCs with SEMA3F KD groups and control CRCs groups, the result was consistent with IL-6 mRNA(p<0.01;Figure 5B). When we silenced the IL-6 expression of the two groups by siRNA in hypoxia, the PPAR activity of hLECs which co-culture in control CRCs groups had no changes, conversely, its activity in the SEMA3F KD-CRCs group was significantly reduced to the control groups level (p<0.01;Figure 5C). Furthermore, when SEMA3F KD-CRCs groups loss expression of IL-6, the hLECs tubulogenesis was significantly decreased in hypoxia(Figure 5D). These data showed that hypoxia conditions maybe promote secretion of IL-6 in SEMA3F-deficient CRCs, and enable hLECs mitochondria lipid oxidation to gain energy. Therefore, we measured the change of P-AMPKα and PGC1α protein level in hLECs co-culture with SEMA3F KD-CRCs when we blocked IL-6 activity with IL-6 antibody in hypoxia.The results showed P-AMPKα and PGC1α protein level of hLECs were both decreased in SEMA3F KD groups (Figure 5E). Together, these data suggested that at the hypoxia, SEMA3F KD in CRCs could promote the expression of IL-6, and activate PGCI-PPAR signaling pathways of hLECs to switch on mitochondria lipid oxidation reaction.

**Figure 5.**
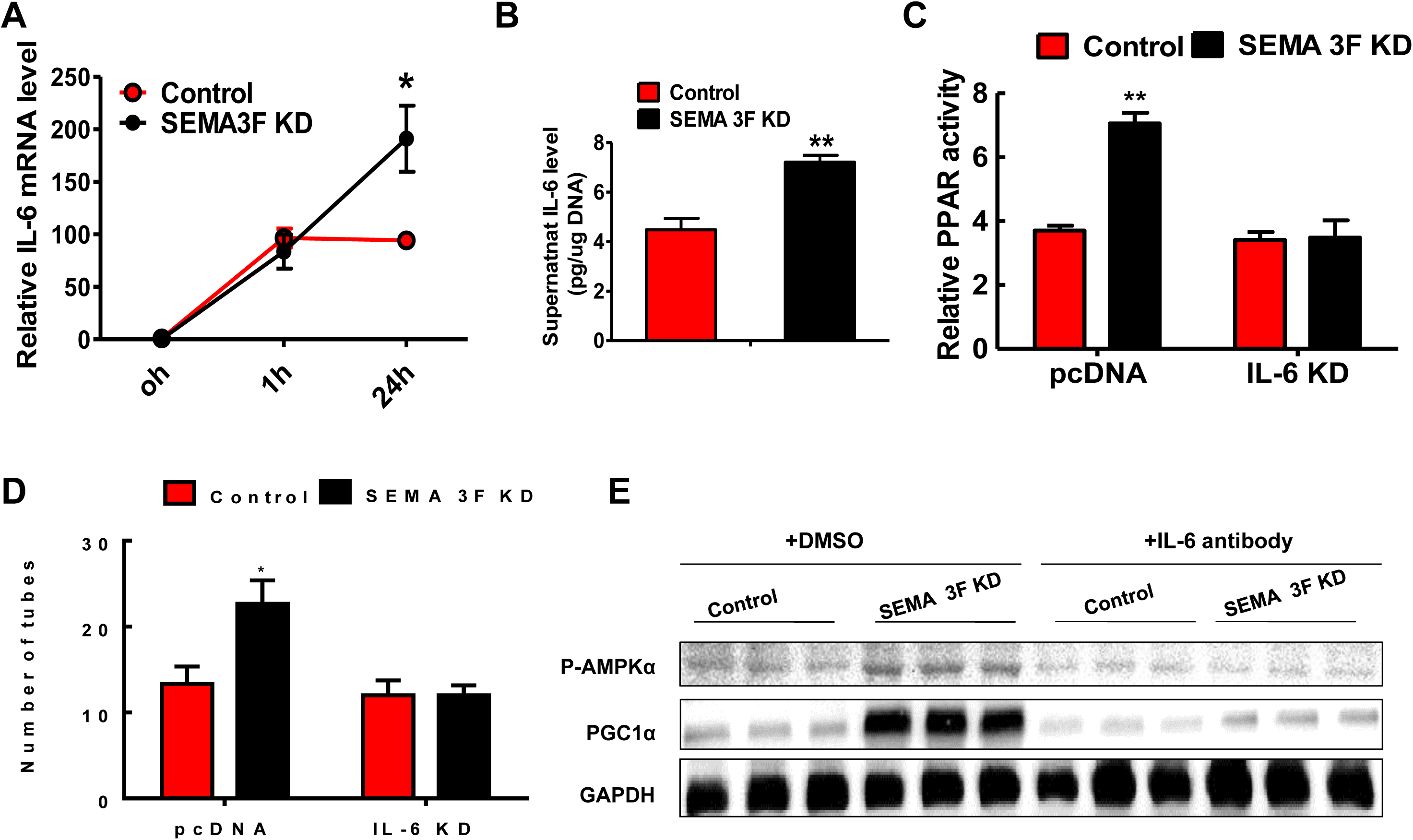
SEMA3F-deficient CRCs under hypoxic conditions promote hLECs PGCI-PPAR lipid oxidation signaling pathway activation by secreting interleukin-6. A, The IL-6 mRNA level of SEMA3F KD-CRCs groups and control groups at the hypoxia. B, The supernatant IL-6 level in CRCs with SEMA3F KD groups and control groups at the hypoxia(**p<0.01). C, Relative PPAR activity of hLECs co-cultured with SEMA3F KD-CRCs group and control group when silencing IL-6 expression of CRCs at the hypoxia. D, Number of tubes sprout from hLECs which co-culture with CRCs in SEMA3F KD groups and control groups at the hypoxia when silenced the IL-6 expression of CRCs. E, Changes of P-AMPKα and PGC1α protein level with western blots analysis in hLECs co-culture with SEMA 3F expression deletion CRCs groups and control groups after adding IL-6 antibody at the hypoxia.

### 6. The results of big data analysis indicate that the SEMA 3F-deficient CRCs showe an increase in hypoxia signal and enhanced self-glycolytic activity

We used gene microarray to investigate the effects of insufficient SEMA 3F expression in 53 CRCs lines on angiogenesis, epithelial-mesenchymal transition, energy metabolism (such as pyruvate metabolism, tricarboxylic acid cycle, glycolysis, and fatty acid metabolism) and so on. The enrichment geneset map was drawn for big data analysis (Figure 6C). The results showed that CRCs lines with SEMA 3F expression deficiency enhanced its hypoxia signal, while induced CRCs epithelial-mesenchymal transition, and promoted angiogenesis ability in colon cancer cell lines. It indicated that the loss of SEMA3F expression could enhance the invasion and metastasis ability of CRCs lines (Figure 6A). Interestingly, the glycolysis of their own was increased, but the aerobic oxidation in mitochondrial was weakened, meanwhile, the fatty acid metabolism of CRCs was also significantly reduced (Figure 6B). Under the hypoxia, the glycolysis of CRCs with SEMA 3F KD was enhanced and consumed a large amount of sugar. It led to insufficient glucose for energy metabolism required for lymphangiogenesis and lymphatic endothelial cell proliferation under hypoxia, this may be the reason why lymphatic endothelial cells take other metabolism to obtain energy. This could explain why the lipid oxidation of mitochondrial was increased while the glycolysis was decreased appeared at lymphatic endothelial cells co-culture with SEMA 3F knockout CRCs.

**Figure 6.**
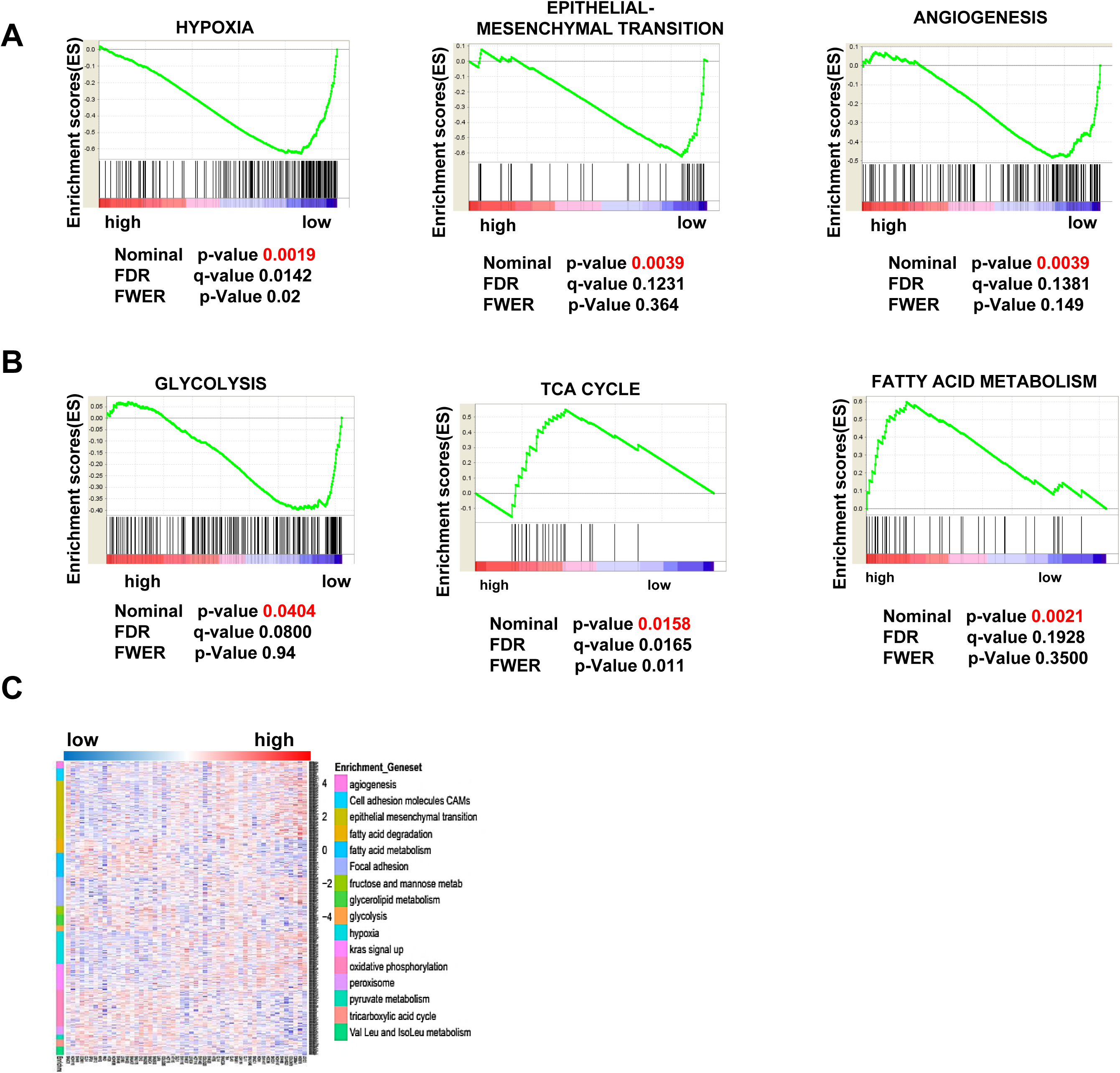
The results of big data analysis showed that the SEMA 3F knockout CRCs increased the hypoxia signal and increased their own glycolysis activity. A. SEMA 3F expression deficiency of 53 CRCs lines enhanced its hypoxia signal, while induce CRCs epithelial mesenchymal transition, and promote angiogenesis. B, SEMA 3F expression deficiency of 53 CRCs lines enhanced its glycolysis,but reduced its fatty acid metabolism and TCA cycle. C, The enrichment geneset in 53 CRCs lines when SEMA 3F expression was deficient.

## Discussion

The tumor metastatic process is a complex multistep process and lymphangiogenesis is the important steps to form distal secondary tumors (Royston *et al*,2009).The initial steps of lymphangiogenesis process that results in cancer cell induction lymphatic endothelial cells proliferation and capillary lymphatic sprouting to elongation of the branch from tumor extravasation into the distal organs (Padera *et al*,2016). Although our understanding of the biological events that energy supply contribute to the lymphatic endothelial cells proliferation is the first event of lymphangiogenesis process over the last decades, we still know very little about the molecular process that includes energy metabolism during lymphangiogenesis, particularly, the hLECs proliferation need energy supply in hypoxemia. In this study, we demonstrated that SEMA3F knockdown in CRCs significantly promoted hLECs migration and tubulogenesis capacity in hypoxia, and activation the lipid metabolism via suppressing aerobic glycolysis of hLECs. To our knowledge, this is the first report demonstrating a molecular event about energy metabolism for the tumor-associated lymphangiogenesis.

It is well known that microenvironment of cancer is often in hypoxic, it induces cancer cell angiogenesis and lymphangiogenesis, the cancer cell proliferation need energy supply too. However, what is the source of energy for these LECs proliferation is not completely clear in hypoxia. In this study, when we slience SEMA3F of CRCs significantly promotes hLECs migration and tubulogenesis capacity, and hLECs proliferation in hypoxia. Interestingly, we have not found the evidence of aerobic glycolysis when hLECs proliferation is induced by CRCs in hypoxia. In contrast, we observe that triglycerides synthesis and lipolysis level in mitochondria of hLECs are significantly elevated. Our results suggest CRCs with SEMA3F KD induce lymphangiogenesis and activate the lipid metabolism in hypoxia. Within the tumor microenvironment, cancer cells consume large amounts of glucose by aerobic glycolysis resulting in hypoxia and hypoglycemia, LECs proliferation experience hypoglycemia and compete for nutrients, and facilitate their switch toward FA catabolism, and preserve their energy supply.

Why the energy supply in hLECs was not come from glycolysis but by lipid metabolism, at the hypoxia condition? However, the energy supply of tumor cell was via glycolysis in hypoxia. We have investigated the effects of insufficient SEMA 3F expression in 53 CRCs lines on angiogenesis, epithelial-mesenchymal transition, energy metabolism and so on, and using large data to analynze it. It indicated that the glycolysis of CRCs own was increased, but the aerobic oxidation in mitochondrial was decreased, meanwhile, the fatty acid metabolism of CRCs was also significantly reduced in hypoxia. SEMA 3F expression deletion CRCs glycolysis wa enhanced and consume a large amount of sugar, it lead to glucose (Glc) depletion within the TME. This metabolic stress in TME led metabolism of LECs switch toward to FAO to gain energy. The metabolic stress of TME also led the lymphatic capillary of tumor to collecting excess fluid and macromolecules from capillary beds almost have not sugar. Why lymphangiogenesis not apply glycolysis metabolism but switch toward to FAO and consume more oxygen? It indicate metabolism of lymphangiogenesis not only for energy supply but also for carbon source for nucleotide synthesis and amino acid synthesis (Schoors *et al*,2015; Zhang, *et al*,2017).This is why the energy supply of hLECs during tumor lymphangiogenesis in hypoxia is lipid metabolism. In recent years, the metabolic reprogramming about T cell in TME had been reported (Zhang, *et al*,2017; Pacella *et al*,2018). T cell enhanced PPAR-α signaling as a fuel switch to promote fatty acid uptakes and utilization. Consistently, in this study showed that CRCs with SEMA3F deficiency not only enhanced hLECs proliferation in hypoxia, but also reprogram LECs energy metabolism. Our data had shown phosphorylation of PGC-1a mediated by AMPK, PGC-1a functions as a coactivator to induce PPARs, such as PPARα and PPARγ, and stimulation of mitochondriogenesis and activation of mitochondrial FAO. It suggested the energy supply of tumor-associated lymphangiogenesis was lipid metabolism and OXPHOS through activation the PGCI-PPAR signaling pathway in hypoxia. In hypoxia TME, the hallmark of cancer cell metabolism is aerobic glycolysis (Warburg, 1956), hLECs facilitate their energy supply switch toward to FA catabolism. however, the more O_2_ amount is consumed through FAO than aerobic glycolysis, it maybe supply other nutrients for LECs proliferation, such as FAO provided acetyl coenzyme A (acetylCoA) to help it deoxyribonucleotide (dNTP) synthesis, it is an anaplerotic substrate for proliferation of LECs (Schoors *et al*,2015).

The past research had shown that IL-6 colud promote the lipid oxidation by raising PPAR in fatty livers (Hong *et al*, 2004; Awazawa *et al*, 2011). In this study, we found the CRCs with SEMA 3F KD could increase its IL-6 secretion, the IL-6 enhanced the PPAR expression of hLECs to drive its lipid metabolism in hypoxia. Wong et al reported that the transcription factor PROX1 upregulated CPT1A and VEGFR3 expression, VEGFR3 ligand VEGF-C elevated FAO in LECs, LECs use fatty acid β-oxidation to proliferate to induce lymphangiogenesis (Wong *et al*,2017). Our data shown that, in the cancer hypoxia microenviroment, CRCs with SEMA 3F KD could promote the secretion of IL-6, and drived mitochondria oxidation reaction in the hLECs by PGCI-PPAR signaling pathways, and not dependent VEGF-C/VEGFR3 signaling. Our findings revealed for the first time a novel role and the underlying mechanism of SEMA3F deficient CRCs involved the energy metabolism with hLECs in hypoxia, and induced tumor lymphangiogenesis of CRCs to promoting lymphatic metastasis.

In conclusion, in this study, our data firstly identified the hypoxia microenviroment promoted tumor lymphangiogenesis and the lipid metabolism was its major energy source when hLECs was proliferation during lymphangiogenesis, however, the glycometabolism of hLECs is suppressed. Our study we also first revealed CRCs secretion IL-6 as a driver, enhanced the expression of PPARa and PGC-1a in hLECs to mediate its the lipid metabolism in hypoxia. When the SEMA3F expression is deficiency in the tumor, the lymphangiogenesis and tumor cell IL-6 secretion and the lipid metabolism are further enhanced. It suggested SEMA3F inhibited tumor lymphangiogenesis not only via NRP2 signaling but also via inhibition LECs energy supply, such as inhibition it’s lipid metabolism.

## Materials and methods

### Isolation of endothelial cells from human tumors

0.5–3g pieces of fresh CRC specimen were cut from the border of the malignancy to be used for the isolation of tumor lymphatic endothelial cells (LEC). The specimen was washed by transferring it using sterile forceps from a 50mL falcon with fresh ice-cold 1× HBSS to a new falcon with 1× HBSS four times consecutively. The specimen was removed from the falcon tube and was placed in a 10-cm cell culture dish to be minced into approximately 10-mm^3^ pieces using a fresh, sterile scalpel. Minced tissue were put into a 15-mL falcon tube filled with 3mL EBM-2MV (Lonza, Cologne, Germany) supplemented with 0.5% FBS. Collagenase II (50mL enzyme per 0.1g tissue; 17,100U/g) was pipetted into each falcon tube, and EBM2-MVwas added up to a total volume of 5mL for each falcon.The falcon tubes were then put in the Dynal sample mixer (Invitrogen, Karlsruhe, Germany) at 37°C for 1h at 5% CO_2_ with the lowest speed available. A 100mm cell strainer (BD Biosciences) was put on a 50mL falcon tube and the digested tissue was poured through the strainer. The filter was washed from the inside and outside using 3mL EBM-2MV each time, and was centrifuged with 500×g for 5min at 20°C to discard the supernatant. The cell pellet was resuspended in 5mL EBM-2-MV, and then was cultivated until 70–80%c onfluence in a T-25 cell culture flask precoated with 1.5% gelatin for at least 2h. The isolated cells can be positively selected for VEGFR3 by MACS (mouse-anti-humanVEGFR3 antibody was purchased from Chemicon, and was ligated to the MACS obtained from Pierce) after the cultures reach 70–80% confluence. MACS selection was repeated until all nonendothelial cells were removed. Contamination with nonendothelial cells was analyzed by staining an aliquot of the cells for VEGFR3, and subsequent immunocytochemical or FACS analysis. The first confluent T-25 flask of pure LECs is designated as passage 0. One passage is defined by a split ratio of 1:4.

### Cell culture

The human colon cancer cell lines LS147T were obtained from the American Type Culture Collection (ATCC, Manassas, VA), and maintained in L-15(Invitrogen Corp.) supplemented with 10% fetal bovine serum at 37 °C under 5% CO_2_. Since both LS147T and HT-29 cell lines expressed SEMA3F, LS147T was less differentiated than HT-29 and was more prone to lymphatic metastasis. Therefore, we selected LS147T cell line as the research object (Supplementary Figure1 A and B).Freshly isolated lymphatic endothelial cells were cultured in ATCC-formulated of F-12K Medium with 0.1mg/mL heparin, 0.03–0.05mg/mL ECGS, supplemented with fetal bovine serum at 37°C under 5% CO_2_. Cells were cultured either in normoxia, in a humidified, 5% CO_2_- and 20% O_2_-containing atmosphere or in hypoxic conditions, which were generated in a humidified hypoxic incubator with 1% O_2_, 5% CO_2_ and balanced N_2_ content (94% N_2_, 5% CO_2_, 1% O_2_).

### Antibodies and reagents

A Mouse monoclonal anti-B-cell lymphoma-2 (Bcl-2) (Catalog #05-729), a Rabbit polyclonal anti- SEMA3F (Catalog#AB5471P), a Rabbit polyclonal anti-HSL(Catalog # ABE204), rabbit polyclonal anti-GPAT1 (ABS764) were a Chemicon International product (Temecula, USA); Monoclonal anti-AMPK and ACC Antibody Sampler Kit Antibodies for AMPK-α, (Catalog #: 9957), Glycolysis Antibody Sampler Kit (Catalog #: 8337) including Hexokinase I (C35C4) Rabbit mAb #2024, Hexokinase II (C64G5) Rabbit mAb #2867, Pyruvate kinase (PKM1/2) (C103A3) Rabbit mAb #3190, Pyruvate Dehydrogenase (C54G1) Rabbit mAb #3205, Lactate dehydrogenase (LDHA) (C4B5) Rabbit mAb #3582, PKM2 (D78A4) XP® Rabbit mAb #4053, Phosphofructokinase (PFKP) (D4B2) Rabbit mAb #8164, rabbit anti-cleaved9(mAb#7237), anti-Caspase3(mAb#9665),HRP-conjugated secondary antibodies and a monoclonal anti-GAPDH (Catalog #2118) antibody for Western blotting were purchased from Cell Signalling Technology (Beverly, MA, USA). A rabbit polyclonal antibody against human Glut1 (H-43) (Catalog sc-7903), a mouse monoclonal antibody against chicken β-actin (C-4) (Catalog sc-47778, known to recognize mouse, human, rat, and chicken β-actin), mouse anti-PPAR-α (mAb sc-130640) and mouse anti-PPAR-γ(mAb sc-74517), rabbit polyclonal Anti-PGC1(sc-13067), mous anti-Cyclin D1 (mAb sc-70899) and mouse anti-Cyclin D2(mAb sc-166288), PLC, mouse anti-DGAT (mAb sc-271934) were purchased from Santa Cruz Biotechnology, Inc. A mouse anti-human PARP (mAb#551025) was purchased from BDBio sciences (SanDiego,CA). A polyclonal rabbit anti-AGPAT (SAB2700797) and a monoacylglycerol lipase An MAGL rabbit polyclonal antibody (Cat. 011994) were purchased from Sigma-Aldrich. A rabbit anti-ATGL (mAb ab207799) was purchased from Abcam. A mouse monoclonal anti-PAPase (AM32058PU-N) was purchased from Acris. A rabbit polyclonal antibody Anti-CGI-58 (#PAB12500, Abnova) was purchased from Abnova. A rabbit polyclonal antibody against human DAGL (diluted 1∶2000, cat. DGLa-Rb-Af380) was purchased from Frontier Science. The GW9662, antagonist of PPAR-γ was purchased from Cayman Chemicals (St Louis, MO, U.S.A.)

### Transfection plasmid information and establishment of stable cell lines

The SEMA3F expression vector pSectag-SEMA3F was kindly provided by Dr. David Ginty, and used as described previously. (Wu *et al*,2011) SEMA3F RNAi expression vectors and control plasmids were obtained and used as previously described (Wu *et al*,2011). Small interference RNAs (siRNAs) against human IL-6 (#sc-39627), AMPKα1(#sc-29673) and PGC1α(#sc-38885) were obtained from Santa Cruz Biotechnology in deprotected and desalted form. All resultant constructs were verified by DNA sequencing, and then transfected into target cells with lipofectamineTM 2000 transfection reagent (Invitrogen, Carlsbad, CA, USA). Transfected cells were enriched by selection for 1 week with antibiotics selection.

### Protein extraction and western blotting

Cell lysates were prepared with M-PER Mammalian Protein Extraction Reagent (PIERCE, PA, USA). A total of 30 μg of lysate proteins were separated by SDS-PAGE after heat denaturation, transferred onto PVDF membranes, and incubated with 5% non-fat milk dissolved in PBS-Tween 20 solution for 1 h, followed by incubation with a primary antibody overnight at 4 °C. After washing, the membranes were incubated with an appropriate HRP-conjugated secondary antibody, and then developed with enhanced chemiluminescence (ECL) detection reagents (Amersham Pharmacia Biosciences).

### Cytokine array/ELISA assay

IL-6 concentrations in the cell supernatant were were detected utilizing mouse IL -6 ELISA kit t (A015171517) purchased from GenScript Biological Technology Co.Ltd. (New Jersey, United States,)and according to the kit instructions.

### Immunofluorescence microscopy studies

Samples for immunofluorescence staining were fixed in ice-acetone for 20 min, washed with PBS 3 times for 5 min each, and incubated for 30 min at room temperature in a protein-blocking solution. The sections were incubated with the primary antibodies for 1 h at 37 °C and then at 4 °C overnight. After washing, the sections were incubated at 37 °C for 1 h with appropriate secondary antibodies, including FITC-conjugated goat anti-rabbit IgG (1:50, Santa Cruz), FITC-conjugated goat anti-mouse IgG (1:50, Santa Cruz), or TRITC-conjugated goat anti-mouse IgG (1:50, Beyotime, China). The sections were counterstained with Hoechst 33258 to reveal cell nuclei.

### Transwell assay

The migration ability of cells was assessed using Transwell chambers with polycarbonate membrane filters with 24-well inserts (6.5 mm diameter and 8 µm pore size) (Corning Life Sciences, Corning, NY, USA). The membrane filters were coated with 1.5 mg/ml Matrigel™ (BD Biosciences, Franklin Lakes, NJ, USA) before use. A total of 5000 cells in 150 µl of McCoy’s 5A-Modified Medium with antibiotics but without serum were seeded onto the upper chamber. The lower chamber was filled with 600 µl McCoy’s 5A-Modified Medium supplemented with antibiotics and 30% FBS. The medium in both chambers was changed once daily. After culture for 48 h, following removal of the non-migratory cells from the upper surface of the filter using a Q-tip, the migrated cells were fixed with cooled-acetone (4°C), and then stained with crystal violet solution (Invitrogen) and counted under ten different low-power (100×) microscopic fields. The cell border was verified by switching to the high-power objective lens (400×) during counting.

### Tubulogenesis assay

LECs were cultured in ECM supplemented with 20% FBS for 12h, and then digested with trypsin /EDTA to prepare for cell suspension. The cell suspension(1.0×10^6^/ mL) was seeded onto a 24-well plate coated with 0.5mL of 4% matrigel, and was then co-cultured with or without CRC cells at 37 °C, 5% CO_2_. Cells were photographed every 2 days and the quantities of the branches of tube-like structures were counted (one branch as one tube).

### Flow cytometry of apoptosis and cell cycle assay

hLECs apoptosis was assessed by flow cytometry of Annexin V/PI (Sigma) staining. After harvesting, the cells were washed twice with PBS and re-suspended in 200 µl of 1x Annexin binding buffer. 5µl Annexin V-FITC and 5 µl PI (propidium iodide) were then added to the cell suspension, and incubated at 37°C for 15 min. The stained cells were analyzed with FACS system (FACS Aria, BD Bioscience). hLECs cycle distribution was analyzed by flow cytometry using propidium iodide (PI) DNA staining (Visagie MH and Joubert AM, 2011). The cells were plated in a six-well plate at a density of 2 × 10^5^ cells per well, and the next day, the cells were treated with fixed concentrations of the fractions and cisplatin. After 24 h, the cells were disrupted and incubated with 40 µg mL^−1^ of PI (Cycle Test Plus BD solution, # 340242) for 10 min at 37 °C, 5% CO_2_, as instructed by the manufacturer. Analysis of the PI-labeled cells was performed by a flow cytometer and the cell cycle phases’ distribution was determined as at least 20,000 cells.

### The cell triglyceride level assay

Harvest the amount of hLECs necessary for each assay (initial recommend = 1 x 10^7^ cells). Wash cells with cold PBS two times. Resuspend and homogenize samples in 1 mL of 5% NP-40/ddH2O solution. Slowly heat the samples to 80 – 100°C in a water bath for 2 – 5 minutes or until the NP-40 solution becomes cloudy, then cool down to room temperature. Repeat previous step to solubilize all triglycerides.Add 2 µL Lipase to Standard and Sample wells. Add 2 µL Triglyceride Assay Buffer (Abcam #ab65336; Abcam, Cambridge, UK) to Sample Background Control wells (do not add Lipase to these samples). Mix and incubate for 20 minutes at room temperature to convert triglyceride to glycerol and fatty acid.Add 50 µL of Reaction Mix into each standard, sample, and background control wells. Mix and incubate at room temperature for 60 minutes protected from light. Measure output on a microplate reader at OD 570 nm for colorimetric assay or at Ex/Em = 535/587 nm for fluorometric assay. The reaction is stable for at least 2 hours.

### The cell total cholesterol (TC) and free cholesterol Level Assay

The TC and free cholesterol content of hLECs were measured colorimetrically using a Cholesterol assay kit according to the manufacturer’s protocol (#ab65359; Abcam, Cambridge, UK). Harvest the amount of hLECs necessary for each assay (initial recommend = 1 x 10^7^ cells). Wash cells Cells were washed in ice-cold saline two times.Cholesterol and cholesteryl ester contents in cells were measured using cholesterol assay buffer after extracted with chloroform: Isopropanol: NP-40 (7:11:0.1). All samples were incubated with the cholesterol assay reaction buffer at 37°C for 60 min. The absorbance was measured at 570 nm.

### The cell phospholipidosis level assay

the phospholipidosis assay was conducted using the hLECs.The LYSO-ID Red cytotoxicity kit (Enzo Life Sciences, Farmingdale, NY, USA) was used for the PLD assay. The LYSO-ID Red dye is a fluorescence reagent that accumulates in lysosomes. Harvest the amount of hLECs necessary for each assay (initial recommend = 1 x 10^7^ cells). Wash cells with cold PBS two times.The cells were stained with LYSO-ID Red dye according to the manufacturer’s instructions. The fluorescence intensity(535 nm excitation, 670 nm emission) was measured using a microplate reader (Infinite F500, Tecan Japan,Kanagawa, Japan).

### Preparation of 3-Bromopyruvatic acid solution

The 3-Bromopyruvatic Acid (3-BrPA) powder (Sigma) was dissolved in PBS. The pH value of this solution was adjusted to 7.0 with NaHCO3. This neutralized 3-BrPA solution was then sterilized with a 0.22 µm filter (Millipore), and used immediately.

### Lipase activity assay

The lipase activity of cells was assessed using the fluorogenic ester substrate 4-methylumbelliferyl heptanoate (MUH) (Sigma). The stock solution of the substrate was prepared by dissolving 1.8 mg of 4-MUH in 1.5 ml methoxyethanol, followed by dilution to 25 ml with distilled water for a final concentration of 1 mM ester. The cells were washed with Tris-buffered saline for 3 times, and then harvested in an ice-cold homogenization buffer [50 mM Tris-HCl (pH 7.4), 250 mM sucrose, 1 mM EDTA]. The enzymatic reaction was initiated by the injection of 60 µl of 2.5 µM MUH in a solution containing 20mM Tris-HCl, pH 8.0 and 1 mM EDTA to 40 µl (100 µg) of the cell homogenate in a 96-well plate (total volume 100 µl). The plate was agitated at room temperature and fluorescence was read with a Fluoroskan Ascent FL Type 374 (Thermo Labsystems) in a kinetic fashion up to 10 min (excitation/emission wavelengths of 355/460 nm). Data were analyzed using GraphPad Prism 5 software.

### Deoxyglucose uptake and lactate assays

For glucose uptake, the cells in 6-well plates were incubated with Krebs–Ringer phosphate buffer supplemented with 1% BSA at 37°C for 30 min. The cells were washed with PBS, and glucose uptake was initiated by adding 2 ml serum-free and glucose-free DMEM containing 2-[1,2-3H(N)]-2-deoxy-D-glucose (DG) (NET328, specificity 8 Ci/mmol) at 1µCi/ml and 10 mM unlabeled-2-DG (Sigma, Catalog #: D-3179) to each sample in the absence and presence of 20 µM cytochalasin B (Sigma Catalog #: C6762), a potent inhibitor of glucose transport. After 5 min incubation, the glucose uptake was terminated by washing the cells rapidly with ice-cold Krebs–Ringer phosphate buffer containing 0.2 mM phloretin (Sigma, Catalog #: P7912). The cells were then solubilized in 500 µl of 0.1% SDS, of which 5 µl was used for the determination of protein concentration, and the rest for liquid scintillation counting after adding 5 ml of Bio-Safe II Cocktail (Fisher, Catalog #: M1-11195). For insulin stimulated-glucose uptake assay, prior to addition of DG, the cells were washed with the insulin-free Stimulation Medium (serum-free and glucose-free DMEM with antibiotics, L-glutamine and 20 mM HEPES), and then incubated with 1 ml of Stimulation Medium supplemented with 0.5 nM human insulin (Sigma) for 5 min. Other steps were identical to those described above for non-stimulated glucose uptake. All experiments were done in triplicate.

### Fatty acid oxidation assay

The incubation medium containing 0.05 µCi/ml [14C]palmitic acid (ARC0172A, specificity 58 mCi/mmol), 0.8 mM unlabeled-oleic acid, and 170 µM fatty acid-free BSA in PBS was prepared before use. The cells were incubated with the incubation medium at 37°C in a 25-cm^2^ Corning cell culture plastic flask whose mouth was tightly covered with a square of lab wipe tissue (Kimwipes, Fisher#: 06-666-A) that was pre-rinsed in 1N NaOH for capture of produced CO_2_. One hour after incubation, the covered wipe tissue was removed for scintillation counting. The cells were then lysed with 500 µl of 0.1% SDS, 5 µl of which was used for protein quantification. Fatty acid oxidation was calculated as radioactivity on the wipe tissue that was normalized to the total amount of cell proteins in the flask.

### Seahorse assays for mitochondria functions

Cell mitochondrial oxygen consumption rates (OCRs) were assayed in 96-well plates by using a XF Cell Mito Stress Test Kit (Cat. #: 101706100, Seahorse Bioscience, U.S.) on the XFe96 Extracellular Flux Analyzer (Seahorse Bioscience, U.S.) according the Manufacturer’s instruction. Basal cellular OCRs were recorded in the absence of any treatment. To record OCRs under metabolic inhibitors or uncouplers, cells were first treated with 2 µg/ml oligomycin to inhibit ATPase. To achieve maximal OCRs, the respiratory chain was uncoupled from oxidative phosphorylation by stepwise titration with 1 µg/ml carbonyl cyanide p-(trifluoromethoxy) phenylhydrazone (FCCP). To completely inhibit the mitochondrial respiratory chain, cells were treated with rotenone (Mito Inhibitor B, a Complex I inhibitor) at 1 µg/ml and antimycin A (Mito Inhibitor A, a Complex III inhibitor) at 1 µg/ml. OCRs were expressed as pmoles/min.

### Staining of mitochondria

Cells were incubated at 37°C and 5% CO_2_ with MitoTracker-Red FM (100 nM) (Invitrogen) for 15 min to stain mitochondria and Hoechst 33258 (2.5 mg/mL) for 15 min to stain nuclei and washed with PBS in between. To stimulate the biogenesis of mitochondria, cells were pretreated with interleukin 4 (IL-4) (Cat. # Z02925-10, GenScript) at a concentration of 5 ng/ml for 24h. Images were taken under an immunofluorescence microscope.

### Data collection

CRC cell expression data from Gene Expression Omnibus (GEO) GSE36133, the submission of CCLE(Cancer Cell Line Encyclopedia) by broad institute. All the CRC cell samples were divided into 2 groups according the median of SEMA3F expression.

### Gene set enrichment analysis

Gene set enrichment analysis(Subramanian *et al*, 2005), was performed using java GSEA Desktop Application (Broad Institute) with the hallmark gene sets (n=50) and KEGG gene sets (n=186) implemented in Molecular Signatures Database (MSigDB 6.0), expression data and phenotype data were formatted following the user guide, samples were permutated with SEMA3F expression level 1000 times.

### Statistical analysis

Data are expressed as Mean ± SEM (Standard Error of the Mean). The pathological scoring data of human specimens were analyzed by 2 biostatisticians in the Department of Statistics, The Third Military Medical University, China. The statistical analysis was performed by one-way ANOVA (when >3 groups) or Students t-test (between two groups) using Graph Pad Prism software. The differences between the values were considered statistically significant when P < 0.05.

## Acknowledgements

We would like to thank the referees and editor, who gave us outstanding advice and helped us build the concepts presented in this work. We would like to thank the referees and editor, who gave us outstanding advice and helped us build the concepts presented in this work. We thank Dr Juanjuan Ou, Department of Oncology, Southwest Hospital, Army Medical University,Chongqing, China,for her excellent technical assistance and help in editing this manuscript.The authors thank Dr Feng Wu, Institute of Pathology of Army Medical University, for help in editing this manuscript.We thank the Centre for Advanced Imaging of Army Medical University, for assistance in confocal laser scanning microscopy imaging. Qi Zhou was supported by the Natural Science Foundation of Chongqing Municipal Science and Technology Commission grant cstc2019jcyj-msxmX0711; Miaomiao Tao is funded by Chongqing Health and Family Planning Committee/ Chongqing Municipal Science and Technology Commission grant ZY201802003.

## Author contributions

Min Chen and Qi Zhou designed the study.Qi Zhou,xiaoyuan Fu and Miaomiao Tao made the figures, and wrote the article.xiaoyuan Fu and Miaomiao Tao performed all of the gene transfection, Western blotting, as well as data analysis. Hongbo Ma and Cancan Wang generated hLECs and cell cluture and gene KO. Yanyan Li, Xiaoqiao Hu and XiuRong Qin performed the glycolysis metabolism assay. Renming Lv and Gengdou Zhou performed the fatty acid oxidation assay. Jun Wang and Meiyu Zhou performed the assays for mitochondria functions. Guofa Xu performed the big data analysis and provided supervision, designed the study, and wrote the article. Zexin Wang provided valuable recommendations, contributing to the study designand reagents.

---

**Supplementary Figure1.**
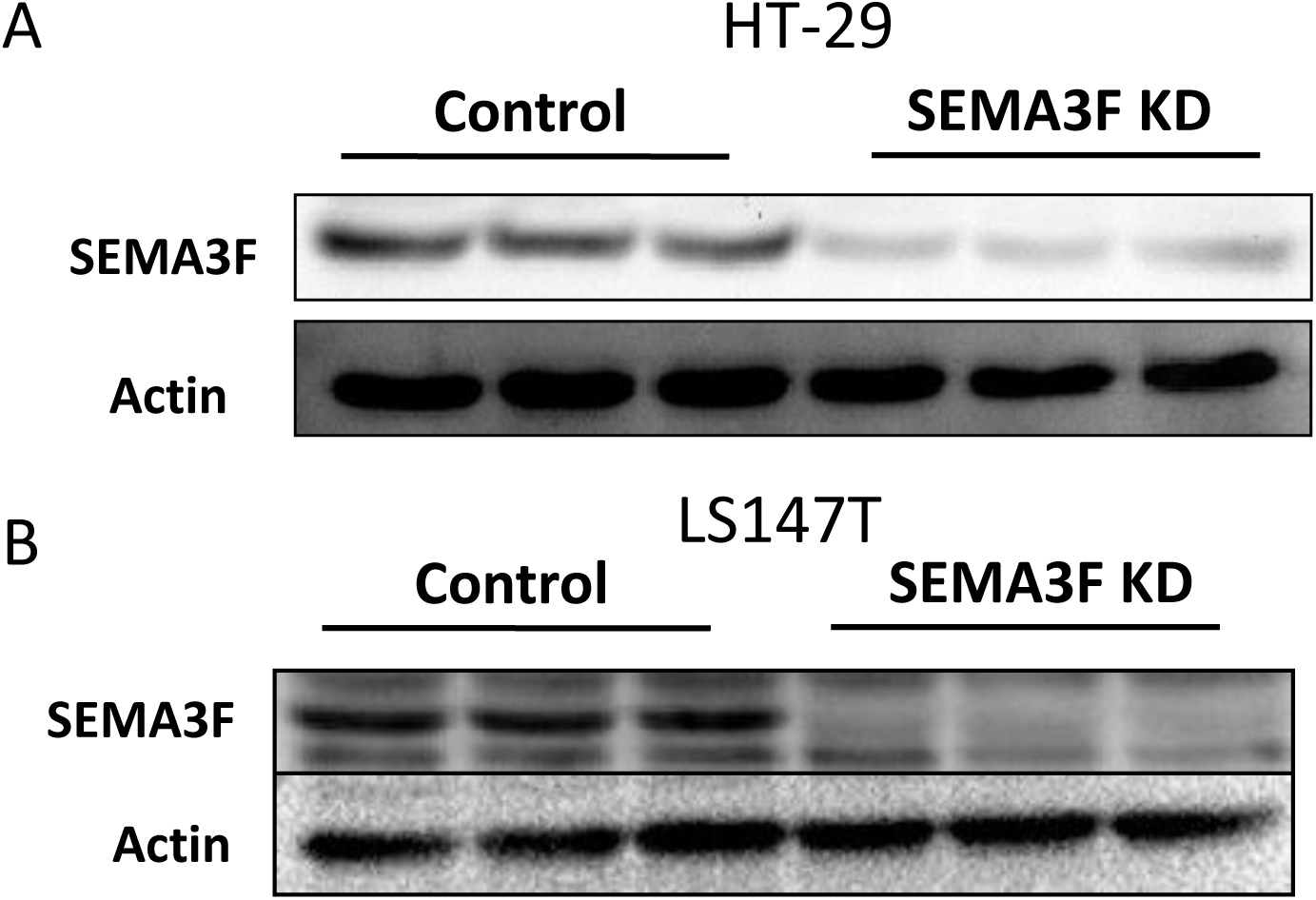
A, HT-29 CRC cell line with SEMA3F exprssion and SEMA3F knocked down. B, LS174T CRC cell line with SEMA3F exprssion and SEMA3F knocked down.

